# PRIGSA2: Improved version of Protein Repeat Identification by Graph Spectral Analysis

**DOI:** 10.1101/803304

**Authors:** Broto Chakrabarty, Nita Parekh

## Abstract

Tandemly repeated structural motifs in proteins form highly stable structural folds and provide multiple binding sites associated with diverse functional roles. The tertiary structure and function of these proteins are determined by the type and copy number of the repeating units. Each repeat type exhibits a unique pattern of intra- and inter-repeat unit interactions that is well-captured by the topological features in the network representation of protein structures. Here we present an improved version of our graph based algorithm, PRIGSA, with structure-based validation and filtering steps incorporated for accurate detection of tandem structural repeats. The algorithm integrates available knowledge on repeat families with *de novo* prediction to detect repeats in single monomer chains as well as in multimeric protein complexes. Three levels of performance evaluation are presented: comparison with state-of-the-art algorithms on benchmark dataset of repeat and non-repeat proteins, accuracy in the detection of members of 13 known repeat families reported in UniProt and execution on the complete Protein Data Bank to show its ability to identify previously uncharacterized proteins. A ∼3-fold increase in the coverage of the members of 13 known families and 3,408 novel uncharacterized structural repeat proteins are identified on executing it on PDB. URL: http://bioinf.iiit.ac.in/PRIGSA2/.

## 1 Introduction

The tandemly repeated structural motifs of length 20-60 residues are most abundant in proteins and assemble to form specific super-secondary structural fold creating favorable protein recognition interfaces (Groves and Barford 1999). These include the super helical structure formed by tandem repetition of anti-parallel helical motifs in α solenoid repeats (eg. Armadillo (ARM), HEAT and Tetratricopeptide repeat (TPR)), horse-shoe structure formed by anti-parallel strand-helix motif in αβ solenoid repeats (eg. LRR repeats), closed propeller structures formed by anti-parallel strands (eg. WD and Kelch repeats) and triple β-spiral structure formed by β-hairpin repeats in trimeric protein complex. The repeat domains interact with DNA, RNA, proteins and small ligands leading to a variety of biological functions and have been implicated in numerous diseases (Pawson and Nash 2003). Members of the same repeat family exhibit varied functions, which may be attributed to the variation in the copy number and sequence similarity between repeating units within a protein (Andrade *et al*. 2001). Low sequence similarity observed between the individual repeating units due to mutations (substitutions, insertions and deletions) accumulated during evolution makes their identification a difficult task. However, accurate detection is necessary for the functional characterization of repeat proteins.

Several algorithms have been developed for the identification of protein repeats at the sequence as well as structure level, capturing distinct features of the repeating units. The sequence based approaches range from methods based on Fourier analysis that capture periodicities in amino acids such as REPPER (Gruber *et al*. 2005) and REPETITA (Marsella *et al*. 2009), to short-string searches such as XSTREAM (Newman and Cooper 2007) and T-REKS (Jorda and Kajava 2009), sequence-alignment based approaches such as RADAR (Heger and Holm 2000), TRUST (Szklarczyk and Heringa 2004) and FAIT (Hrabe *et al*. 2016), and HMM-profile based methods such as HHRepID (Biegert and Söding 2008). The Fourier transform based methods perform poorly in the presence of insertions/deletions within and between the repeat regions which break the periodicity of the amino acids, while the performance of alignment-based methods is affected due to low sequence similarity between the repeating units. The HMM-profile based methods are best suited for the detection of long imperfect repeats but require pre-computed alignments of repeat regions and thus are not suitable for *de novo* detection of novel repeats.

Various approaches have been proposed for the detection of repeats at structure level. These include computationally intensive self-alignment of the protein structure (eg. DAVROS (Murray *et al*. 2004) and OPASS (Shih *et al*. 2006)), and self-alignment of sequence of α characters derived from the backbone dihedral angles (eg. Swelfe (Abraham *et al*. 2008) and ProSTRIP (Sabarinathan *et al*. 2010)). RAPHAEL generates geometric profiles based on C_α_ coordinates and uses them in combination with support vector machine (SVM) to mimic visual interpretation of a manual expert and classifies a protein into solenoid/non-solenoid class (Walsh *et al*. 2012), while graph based approach, ConSole (Hrabe and Godzik 2014), uses a rule based machine learning technique to identify solenoid repeats in proteins. TAPO (TAndem PrOtein detector) (Do Viet *et al*. 2015) considers various structural features such as periodicities of atomic coordinates, strings generated by conformational alphabets, residue contact maps and arrangements of vectors of secondary structure elements to build a prediction model using SVM for identification of structural repeats. ReUPred (Hirsh *et al*. 2016) is another structure based method which predicts repeats by performing iterative structural comparison against a manually refined library of representative repeat units. The methods based on the periodicity of dihedral angles perform poorly in the presence of large insertions/deletions while the scope of learning based methods are limited to the availability of training datasets.

Structural stability of a repeat domain is governed by the packing interactions within a repeat unit and the stacking interactions between repeating units (Main *et al*. 2003). For example, a repeating unit of two anti-parallel helices may form a curved horse-shoe fold as in Ankyrin repeat proteins, or a super helical structure as in TPR repeat proteins, or a closed structure as in protein prenyltransferase subunit beta (PFTB) repeat. The knowledge of different structural repeats reported till date is miniscule of all repetitive structural conformations possible in protein structures. In order to bridge this gap and identify novel previously uncharacterized repeat types, *de novo* methods purely driven by structural properties are required. Here, we present an improved version of our earlier algorithm, PRIGSA (Chakrabarty and Parekh 2014c), that captures inter- and intra-repeat unit interactions typically observed in class III and IV repeats of Kajava’s classification (Kajava 2001, 2012). This is done by analyzing the repetitive pattern in eigenvector centrality profile of the network representation of protein structures. To the best of our knowledge, PRIGSA2 is the only structure based approach which integrates knowledge based identification of repeats along with *de novo* identification. The method extends the coverage of known protein repeat families on one hand, and also enables identification of novel repeat types not yet reported in any of the protein pattern/repeat databases such as Pfam (Finn *et al*. 2016) and PROSITE (Sigrist *et al*. 2013). Further, it is also able to detect structural repeats formed in multimeric states.

## 2 Method

The flowchart of PRIGSA2 algorithm is given in figure 1, with the modifications incorporated in this version marked in ‘red’ color. The algorithm has been tailored to detect Class III and IV repeats (Kajava’s classification) that form elongated and closed structures respectively (size: 5-60 residues) (Kajava 2001). The algorithm comprises 3 major modules: (i) Network Construction Module, (ii) Repeat Identification Module and (iii) Post-processing Module.

**Figure 1.**
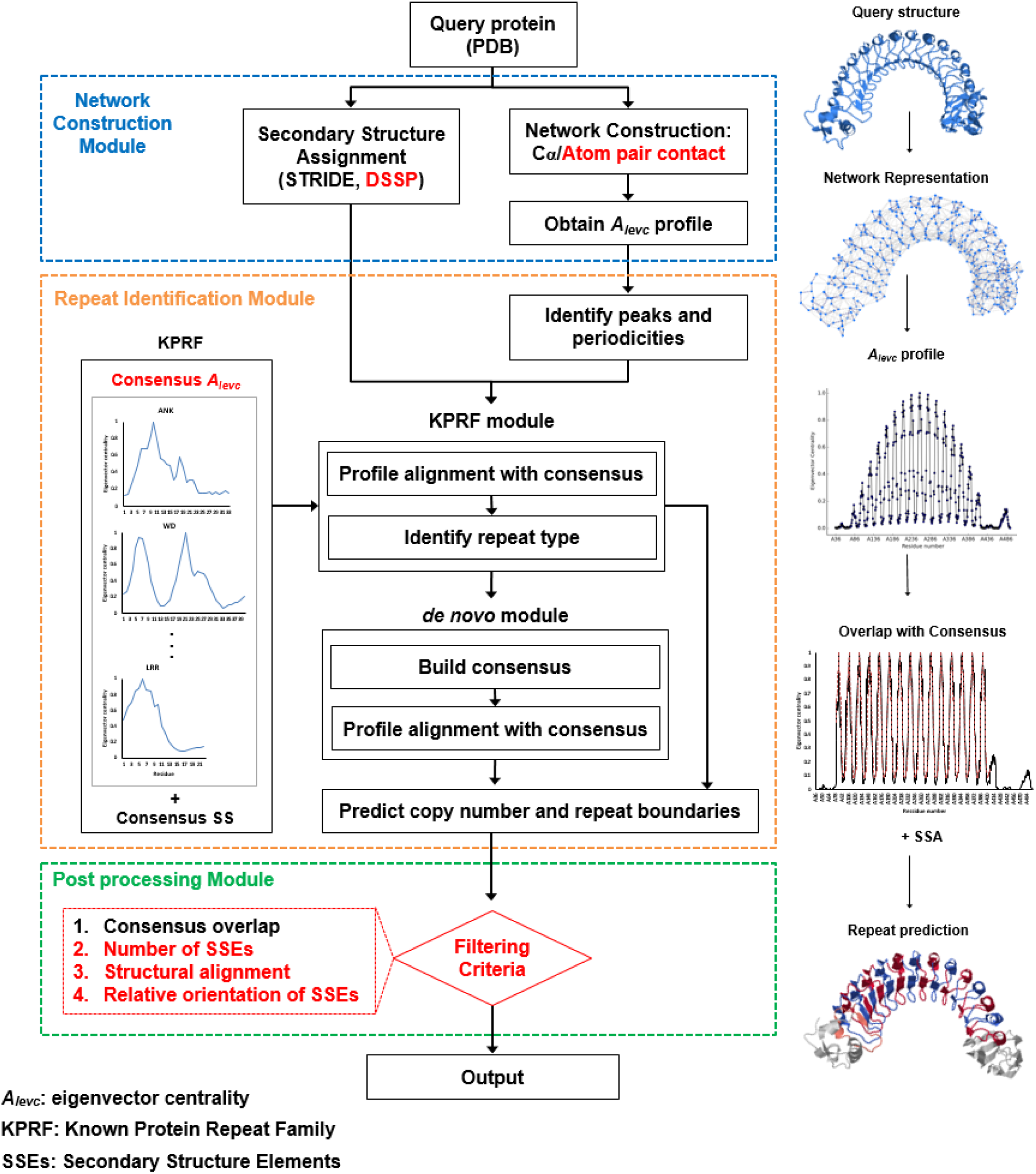
Flowchart showing the steps of the PRIGSA2 algorithm. The modifications in the algorithm with respect to the previous version are marked in red color.

### 2.1 Network Construction Module

The algorithm takes the protein structure file in PDB format as input and constructs a protein contact network (PCN) for the specified chain with Cα atoms as nodes and an edge drawn between them if the Cα-Cα distance ≤ 7 Å. This connectivity information between the residue pairs is represented by *n* × *n* symmetric adjacency matrix, where *n* is the number of nodes (amino acids) in the PCN. The principle eigen spectra of the adjacency matrix (i.e., the vector components corresponding to the largest eigenvalue, *A_levc_*) is known to contain the contribution of each node in the graph, as apart from the connectivity of a node, it also captures the connectivity of its neighbors and their neighbors and so on (Patra and Vishveshwara 2000). Consequently, in the repeat region, the *A_levc_* profile exhibits a similar pattern for each repeating unit (Chakrabarty and Parekh 2014a). This feature (similarity in the *A_levc_* profile) along with secondary structure architecture of the repeat motif is used in the detection of class III and IV repeats (Chakrabarty and Parekh 2014b).

Recently, structural repeats have been observed in multimeric proteins, wherein the structural repeats are observed only in the *k*-meric state but not in monomer chains. For example, for some β-hairpin repeats, stable tandem structural repeats have been observed by forming H-bonds between corresponding residues of multiple chains while no structural repeat is observed in independent folds of monomeric chains (Roche *et al*. 2018). For the detection of this class of structural repeats in *k*-meric states, the atom-pair contact network representation of the protein complex (all the chains taken together) is now provided in PRIGSA2. An atom pair contact network is constructed in this case by considering amino acid residues as nodes and an edge is drawn if the distance between any two atoms of the corresponding residue pair (within or between protein chains) is ≤ *R_c_* ∼5Å (Chakrabarty and Parekh 2016).

### 2.2 Repeat Identification Module

The repeat detection module in PRIGSA2 comprises two sub-modules: (i) knowledge-based, and (ii) *de novo*. As discussed in our previous work (Chakrabarty and Parekh 2014b, c), the eigen spectra of the adjacency matrix, *A_levc_*, and the secondary structure assignment obtained using STRIDE and DSSP databases are inputs to this module. The *A_levc_* profile is analyzed for periodicities in inter-peak distances based on which the length of the repeating units is identified. Next, a consensus *A_levc_* profile is built dynamically or pre-computed consensus profiles for known families (as discussed below) and consensus secondary structure architecture is used for detecting the repeat boundaries and copy number (as discussed in our earlier paper (Chakrabarty and Parekh 2014c)). Since repeat annotation is available for a large number of proteins in UniProt, we utilize this information in PRIGSA2 in our knowledge-based module to improve the prediction accuracy of members of known protein repeat families. Based on available information, we have pre-computed a library of representative *A_levc_* profiles and secondary structure architectures for 13 known repeat families belonging to class III and IV. The repeat families are selected based on their annotation in UniProt, availability of structural information in the repeat region for at least 5 unique UniProt entries of the repeat family, and length of the repeat unit ≤ 60 residues (restricting to Kajava’s class III and IV repeats). Thirteen protein repeat families thus identified are Ankyrin (ANK), Armadillo (ARM), HEAT, Pumilio, protein prenyltransferase subunit alpha (PFTA) family, protein prenyltransferase subunit beta (PFTB) family, Tetratricopeptide repeat (TPR), Leucine rich repeat (LRR), Parallel beta-Helix repeat (PbH1), Kelch, WD, Hemopexin and Low Density Lipoprotein Receptor class B (LDLR-B). These 13 repeat families belong to five structural sub-classes in Kajava’s classification of class III and IV repeats, namely, α solenoid, αβ solenoid, β solenoid, α barrel and β propeller (table 2).

The query protein is first scanned by the Known Protein Repeat Families module (KPRF) to check if it belongs to any one of the 13 repeat families, before going to the computationally intensive *de novo* module. In the KPRF module, information available on known protein repeat families is utilized to (1) validate the prediction of repeat type, (2) extend the coverage by identifying new members of known repeat families, and (3) improve the annotation of the known members (copy number and start/end of the repeat boundaries). In the *de novo* module, periodicity in all-to-all peak distances in the *A_levc_* profile and their frequencies and periodicity in the secondary structure elements are analyzed to identify novel uncharacterized structural repeats in proteins (see (Chakrabarty and Parekh 2014c) for details). Apart from novel repeats, this module also identifies members of known repeat families missed by the KPRF module (other than the 13 KPRFs), and members of repeat families for which no consensus *A_levc_* profile and secondary structure is pre-computed due to limited structural information in PDB.

### 2.3 Enhancements in PRIGSA

#### 2.3.1 Secondary Structure Assignment

In PRIGSA2, secondary structure (SS) information is utilized in two modules: (i) validating the periodicity predicted by analyzing peak-peak distances in the *A_levc_* profile in the repeat identification module, and (ii) validating the predicted repeat boundaries in the post processing module. In the earlier version, the SS assignment was obtained from STRIDE program (Frishman and Argos 1995). We observed that because of inconsistent or no assignment given by the STRIDE program, some known repeat proteins were missed by PRIGSA. To handle this issue, in PRIGSA2 the SS assignment is obtained from both STRIDE and DSSP (Kabsch and Sander 1983); if the algorithm terminates due to an error in STRIDE execution, SS assignment is considered from DSSP. Further, in the post-processing step, a repeat prediction is accepted if SS architecture of the predicted repeat unit (either by STRIDE or DSSP) is in agreement with that of known repeat family or predicted consensus. Though both STRIDE and DSSP show an agreement of more than 95% (Cuff and Barton 1999), the prediction accuracy of PRIGSA2 improved when both STRIDE and DSSP are considered.

#### 2.3.2 Construction of Representative Profiles for Known Protein Repeat Families

In the earlier version of our algorithm, the representative *A_levc_* profile and SS architecture were considered for 8 known repeat families (ANK, ARM, HEAT, Pumilio, TPR, LRR, Kelch and WD). The *A_levc_* profile of an intermediate copy of a designed protein (if available), or the best resolution structure was considered as the ‘representative profile’ for the respective family.

However, it is observed that many members of the family exhibit deviation from the designed/best resolution structure due to variations in the repeat region. Moreover, for some families (e.g., LRR), sub-classes have been reported and a single profile may not suffice in such cases for identifying all valid members of the repeat family. In PRIGSA2, the consensus profiles have been constructed for 13 protein repeat families, given in table S1 (compared to 8 in PRIGSA). Further, an elaborate procedure is now followed in the construction of the consensus *A_levc_* profile(s) as depicted in figure 2. First, the *A_levc_* profile of all repeat copies is extracted for each family based on the annotation of repeat boundaries in UniProt. An all-against-all *A_levc_* profile-profile alignment is performed with periodic boundary conditions by aligning all peaks of the two profiles and a similarity score, *S*, is computed as:

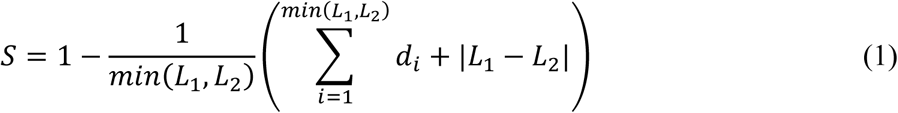

where, *L_1_*, *L_2_* are lengths of the two copies being compared and *d_i_* is the difference in the *A_levc_* values at position *i*. The second term |*L_1_* – *L_2_*| penalizes the difference in the length of the repeat units. Thus, higher values of *S* indicate greater similarity between the repeat motifs. An unweighted network is constructed for each repeat family by considering all the reported repeat copies in UniProt as nodes and an edge drawn between two repeat units if the score between their *A_levc_* profiles is greater than a threshold score, *S_t_*. This network is subjected to Markov Clustering algorithm (MCL) (Enright *et al*. 2002) to identify sub-groups, if any, within each repeat family. The clusters with 30 or more members (repeat units) are considered to build consensus profiles. The *A_levc_* profiles of all members of a cluster are aligned by the principal peak in the profiles and a consensus is constructed by averaging the *A_levc_* values at each position of the aligned profile. This helps in identifying more than one representative profile for a repeat family in case of large variations in the members of the repeat family or sub-classes of the repeat family, as shown in table S1. A representative structural motif is obtained for each cluster (corresponding to each *A_levc_* profile) by carrying out an all-against-all structural alignment of all the members of the cluster and selecting the repeat motif exhibiting minimum average RMSD with all the members of the cluster.

**Figure 2.**
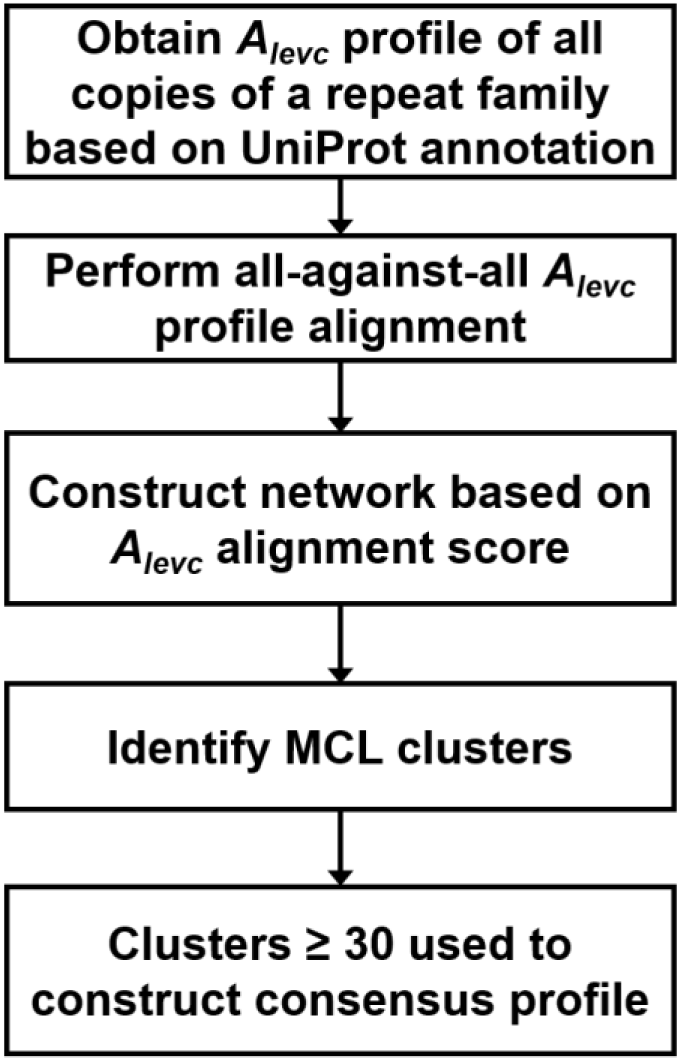
Workflow for the construction of consensus *A_levc_* profile for known protein repeat families (KPRFs).

The consensus profile(s) thus obtained are then used as representative(s) of the respective protein repeat families for predicting the repeat type. Due to large variations in the repeat units from the consensus, some members are missed by the KPRF module of the program. Some of these are detected by the *de novo* module, however, in such cases only the repeat region is reported by PRIGSA2, not the repeat type.

### 2.4 Post-processing Module

To improve the reliability of predictions and reduce false positives, we have incorporated the following additional filtering steps in both KPRF and *de novo* modules. This is done based on the number of conserved secondary structure elements (SSEs) and structure-structure alignment of the predicted repeat copies as discussed below.

#### 2.4.1 Filtering based on Number of Conserved Secondary Structure Elements

The post-processing step in the earlier version of PRIGSA required that at least two copies in the predicted repeat region should have all the SSEs conserved both in number and order of occurrence in accordance with the consensus profile (pre-computed or dynamically obtained). In PRIGSA2, this condition is made more stringent by expecting that over and above at least two copies having all the SSEs, all the remaining copies should contain at least (*n*-1) SSEs, *n* being the no. of SSEs in the consensus secondary structure architecture. This condition has aided in improving the prediction of terminal copies, which are generally incomplete. Since the interaction pattern of the terminal copies is well conserved in closed repeat types (class IV) due to the spatial constraint, this filtering criterion based on SSEs is not necessary (and not carried out in the KPRF module).

#### 2.4.2 Validation of Predicted Repeats

We have incorporated two validation steps in PRIGSA2: (i) structure-structure alignment of predicted repeat copies and (ii) checking orientation of the corresponding secondary structure elements in adjacent repeat units.

#### Structure-structure alignment

The structure-structure alignment of repeat units is carried out by *cealign* algorithm (Shindyalov and Bourne 1998) implementation in PyMOL (The PyMOL Molecular Graphics System). In the KPRF module, structural alignment of each predicted repeat copy with the representative structural motifs of the repeat family is carried out. All copies with RMSD ≤ 3Å are accepted as true predictions. Similarly, in the *de novo* module, an all-against-all pairwise structural alignment between every pair of predicted copies is carried out and the predicted region is accepted to be true if RMSD between two or more pairs is ≤ 3Å. The threshold value of 3Å is arrived at by analyzing all the annotated repeat units from the 13 known repeat families. Pairwise structure-structure alignment of all *vs* all repeat units in each repeat protein annotated in UniProt was carried out and cumulative frequency distribution of the RMSD values obtained. It was observed that ∼ 81% of repeat pairs within a protein had an RMSD ≤ 3Å, and on considering only top two structural alignments in each protein, ∼98% of repeat pairs had an RMSD ≤ 3Å as shown in figure 3 (a) and (b) respectively.

**Figure 3.**
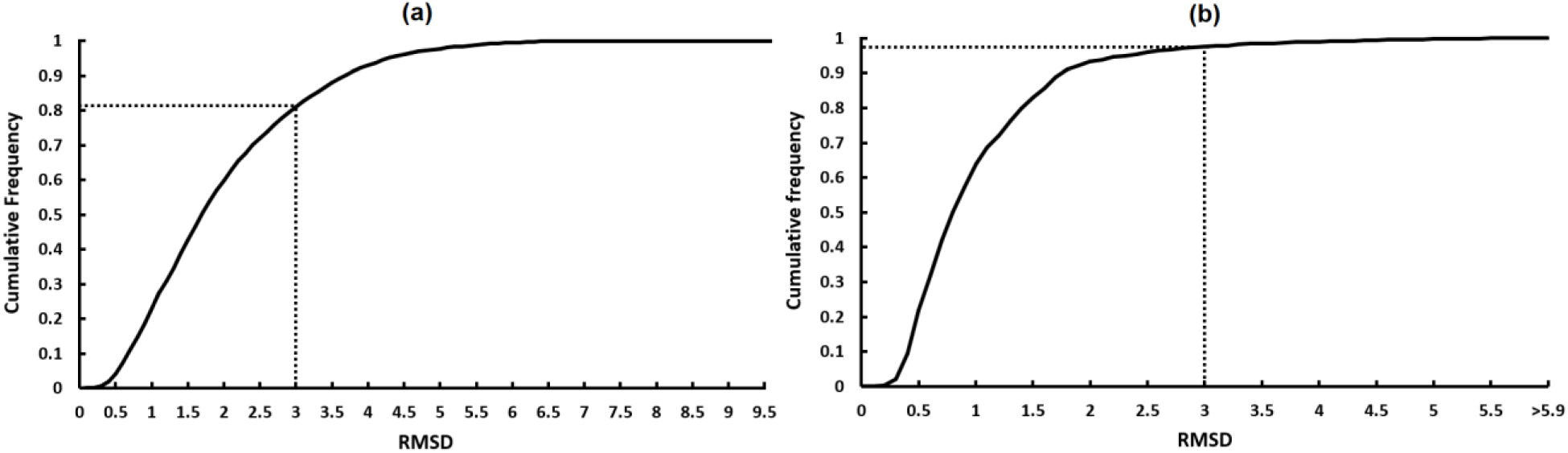
Cumulative frequency distribution of RMSD values obtained from structure-structure alignment of repeat units for 13 known repeat families: (a) all repeat units within a protein (b) two best aligning repeat units within a protein.

#### Relative Orientation of Secondary Structure Elements in Adjacent Repeat Copies

It is observed that the structure-structure alignment of individual repeating units discussed above only ascertains structural similarity at the single motif level, not at the overall tertiary fold of the repeat domain. However, in a tandem repeat region, the overall 3D topology of the repeating structural motif and the relative orientation of secondary structure elements within each motif are well conserved to form a unique overall super-secondary structural fold of the repeat domain. For example, with the earlier version of PRIGSA, Chemotaxis protein CheY (PDB: 1AB5, chain: A), shown in figure 4, was predicted to contain 5 copies of LRR repeat. In this case the average RMSD of all pair-wise structural alignments of the 5 repeat copies is ∼ 2.4Å, suggesting high structural similarity between the repeat copies. However, the typical horse-shoe fold of LRR is not observed in this case, indicating it to be a false prediction of LRR repeat type.

**Figure 4.**
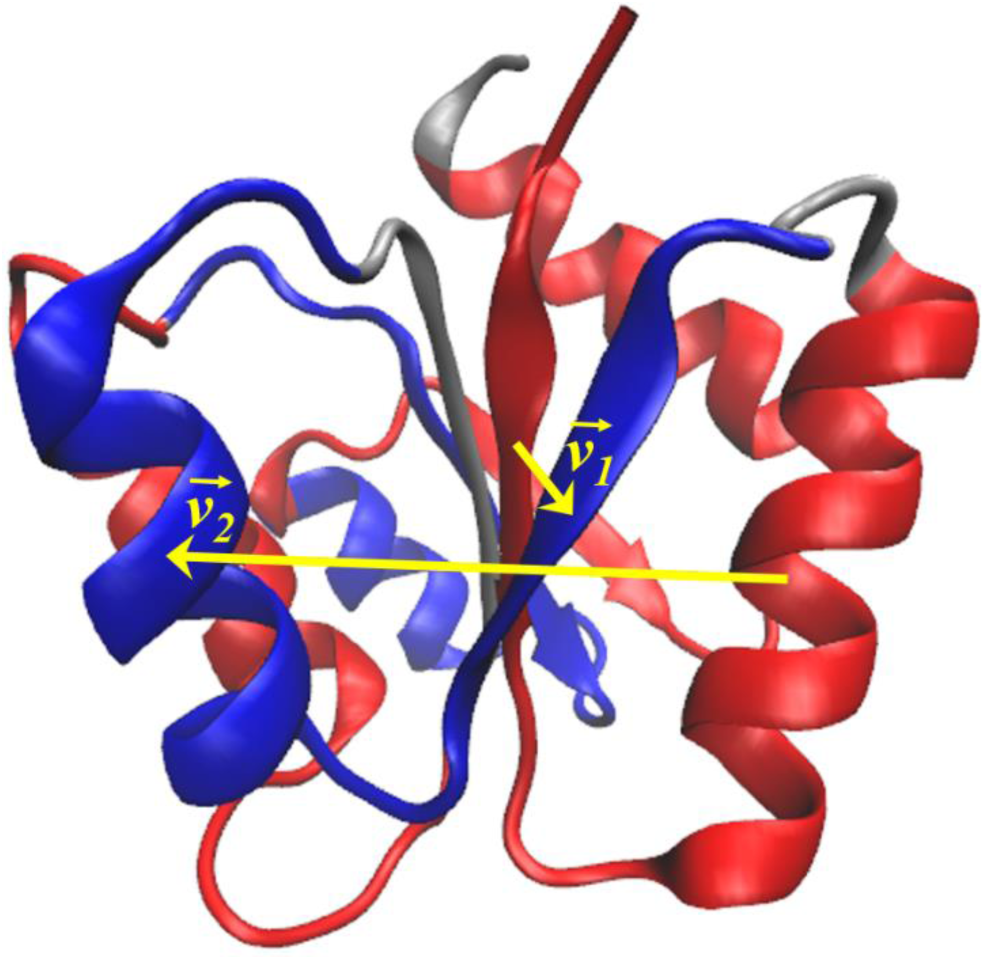
3D structure of Chemotaxis protein CheY (PDB: 1AB5). The alternate repeat copies predicted by PRIGSA1 are shown in red and blue colors respectively.

To address this issue, we have implemented a simple geometric measure to capture the conserved secondary structure orientation in PRIGSA2. In figure 5 (a), a cartoon representation of an ‘elongated’ α/β solenoid repeat with Strand-Helix repeat motif is shown. The vectors, *v_1_* and *v_2_*, joining the central residue of the strands and helices of two adjacent Strand-Helix repeat motifs are observed to be parallel to each other. However, vectors *v_1_* and *v_2_* joining the central residue of the corresponding strands and helices of two adjacent repeat copies in protein 1AB5 in figure 4 are not parallel. In figure 5 (b), ‘closed’ β propeller repeat with four consecutive strands comprising a repeat unit is shown. In this case also, the four vectors (*v_1_*, *v_2_*, *v_3_* and *v_4_*) joining the central residues of corresponding strands of the two adjacent repeat units are observed to be parallel to each other. We exploit this feature of tandem structural repeats to filter out False Positives and improve the prediction accuracy of PRIGSA2. If the angle between the vectors drawn through the central residues of corresponding secondary structural elements in adjacent repeat units is ≤ 30° for at least two pairs of adjacent copies, the prediction is accepted to be true.

**Figure 5.**
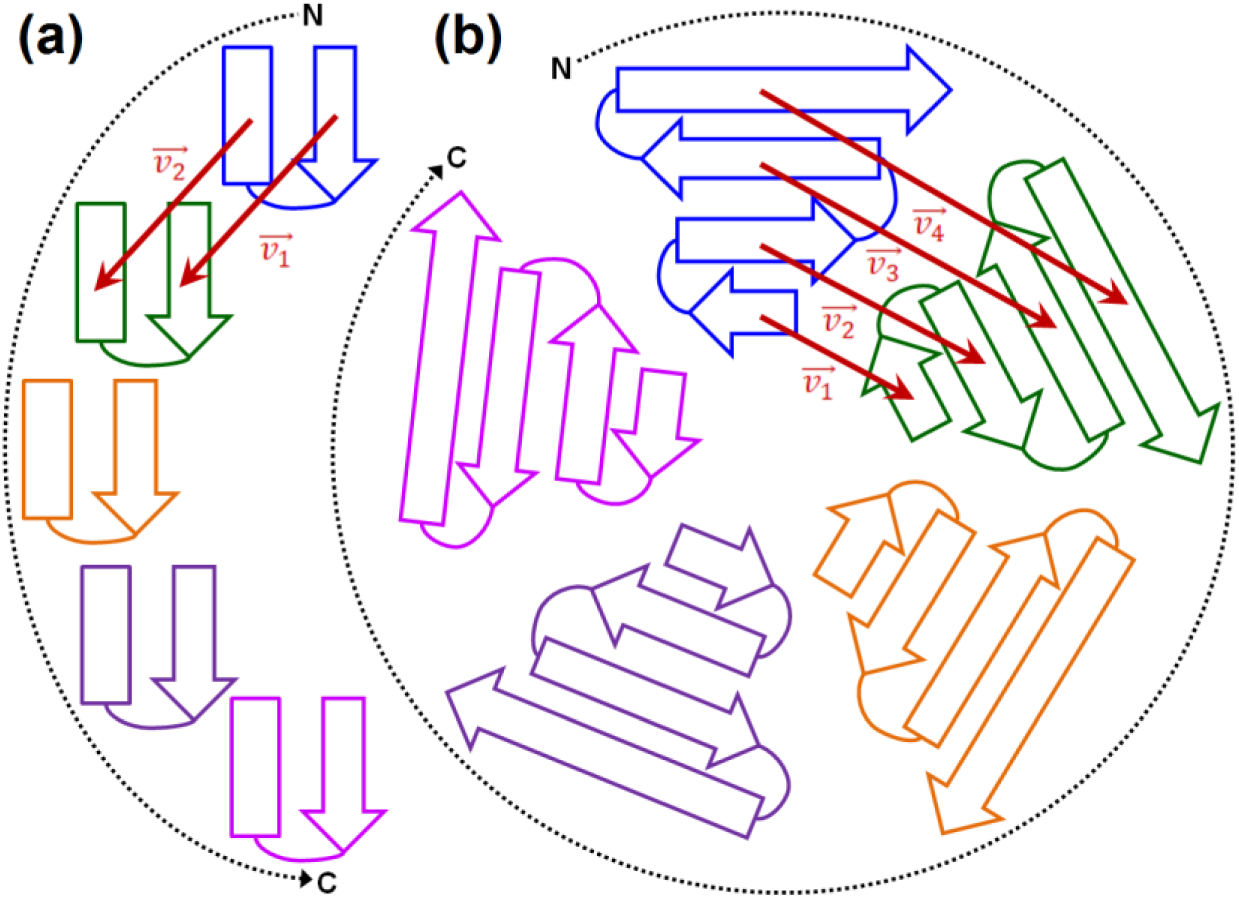
Validation step to check the relative orientation of secondary structure elements. (a) *v_1_* and *v_2_* are the vectors joining the corresponding strand and helix respectively of two consecutive copies of Strand-Helix motif of ‘elongated’ type repeat family forming an horse-shoe structure. (b) *v_1_*, *v_2_, v_3_* and *v_4_* are the vectors joining centers of corresponding strands of a ‘closed’ type repeat family with 4 strands.

### 2.5 Dataset

The performance evaluation of PRIGSA2 algorithm is executed on three datasets: **(i) Benchmark dataset** (Marsella *et al*. 2009): 352 proteins comprising 105 solenoid repeat proteins and 247 non-solenoid proteins. Comparative analysis of the performance of PRIGSA2 with four state-of-the-art algorithms is performed on this dataset. **(ii) Non-redundant dataset of repeat proteins:** 375 proteins comprising members of 13 known protein repeat families (ANK, ARM, HEAT, Pumilio, PFTA, PFTB, TPR, LRR, PbH1, Kelch, WD, Hemopexin and LDLR) belonging to five structural subclasses of Class III and IV in Kajava’s classification (see table 2). This dataset is constructed by considering reviewed entries from UniProt database (release 2017_10) (UniProt Consortium 2015) for which structural data is available in the repeat region with three or more contiguous repeats. The best resolution structure with maximum repeat coverage for each UniProt entry is considered. On this dataset, the prediction accuracy of PRIGSA2 in distinguishing between repeat and non-repeat regions at copy number and residue level is assessed. **(iii) Protein Data Bank (**as on November 3, 2017) (Berman *et al*. 2000): The PRIGSA2 algorithm was executed on the complete PDB containing 4,49,867 chains to highlight the efficacy of *de novo* module in identifying novel, under represented repeat proteins in UniProt and previously uncharacterized repeat proteins.

### 2.6 Implementation details

The PRIGSA2 algorithm is available as a web server which takes the protein structure in PDB format along with the number of chains, chain id(s) and network type to be considered for repeat prediction. The algorithm is implemented in Python and the workflow of the algorithm is shown in figure 1. On submitting a query structure, the PRIGSA2 web server gives the repeat type (if identified as one of the 13 KPRF) or periodicity (for *de novo* predicted repeat), copy number, start/end boundaries of the repeat units and the prediction score for each repeat unit. The 3D structure of the protein is displayed in JSmol applet with alternate copies colored in red and blue colors, as shown in figure 6 for an example protein, 2OMX. The prediction score for individual repeating units is obtained by aligning its *A_levc_* profile with the consensus *A_levc_* profile using equation 1.

**Figure 6.**
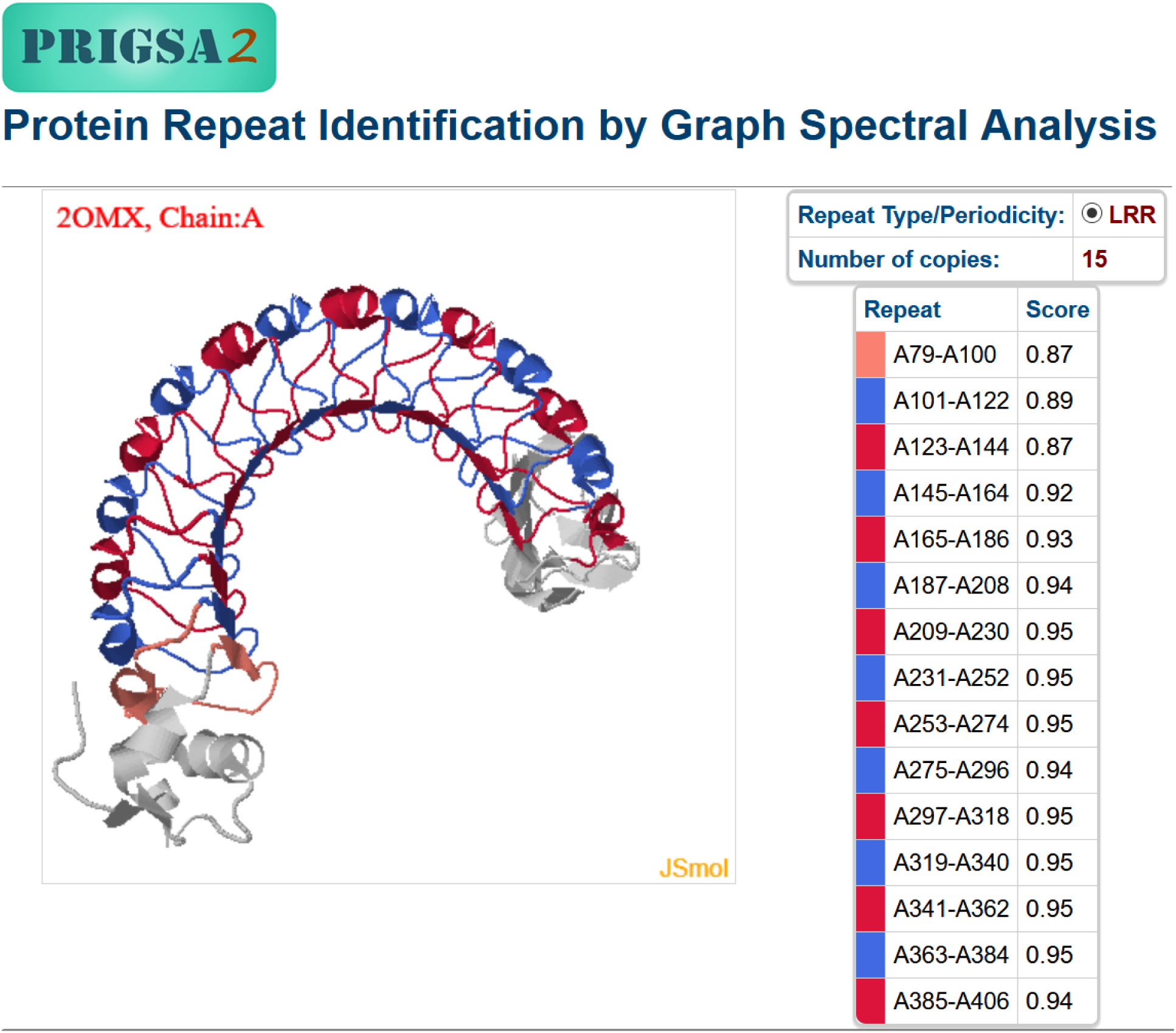
Snapshot of PRIGSA2 output for an example LRR repeat protein, 2OMX (chain A).

## 3 Results

We have developed PRIGSA2 with two objectives: (i) *de novo* detection of structural repeats so as to identify novel uncharacterized repeats, and (ii) provide annotations to uncharacterized members of 13 known protein repeat families (KPRFs) that belong to five subclasses of class III and IV in Kajava’s classification scheme (see table 2). Below we discuss the performance of PRIGSA2 algorithm at three levels.

### 3.1 Benchmarking on Solenoid Repeat Protein Dataset

Performance of PRIGSA2 algorithm is compared with four state-of-the-art repeat detection algorithms, namely, REPETITA (Marsella *et al*. 2009), ConSole (Hrabe and Godzik 2014), TAPO (Do Viet *et al*. 2015) and RepeatsDB-lite (Hirsh *et al*. 2018), and its previous version (PRIGSA (Chakrabarty and Parekh 2014c)). ConSole applies image processing on contact map representation of protein structure and uses a trained SVM to identify repeats, while TAPO ranks the predictions from various sequence and structure based methods on a SVM classifier to identify repeats. The performance is evaluated on a benchmark dataset of 352 proteins comprising 105 solenoid repeat proteins and 247 non-solenoid proteins (Marsella *et al*. 2009) by computing sensitivity (TP/(TP+FN)) and specificity (TN/(TN+FP)) of these approaches, summarized in table 1. The prediction results on the dataset are given in http://bioinf.iiit.ac.in/PRIGSA2/S1_Benchmark_Dataset.xlsx. The objective of this exercise is to test the efficacy of PRIGSA2 in distinguishing repeat proteins from non-repeat proteins. It may be noted from the table that specificity of PRIGSA2 (∼ 0.98) is much higher than all the other methods, indicating its ability to correctly discard non-solenoid proteins. However, the stringent conditions of the algorithm result in relatively lower sensitivity of 0.60 suggesting that some of the true positives are missed. ConSole and TAPO exhibited higher sensitivity (≥ 0.90) but with compromised specificity of ∼0.7. The improved specificity of PRIGSA2 but slightly lower sensitivity compared to its previous version (PRIGSA) is due to the stringent conditions imposed on the conservation of secondary structure elements and structure-based validation steps incorporated in PRIGSA2. The other methods being context based are limited by the datasets used for training, while PRIGSA2 being based on a *de novo* approach, is able to identify any uncharacterized tandem structural repeat.

**Table 1.** Comparison of PRIGSA2 with four structure-based approaches. (The results for algorithms marked by ‘*’ are reproduced from (Chakrabarty and Parekh 2014c))

| Method | Sensitivity | Specificity |
| --- | --- | --- |
| REPETITA* | 0.70 | 0.84 |
| ConSole* | 0.90 | 0.72 |
| TAPO | 0.94 | 0.70 |
| RepeatsDB-Lite | 0.62 | 0.82 |
| PRIGSA* | 0.70 | 0.79 |
| PRIGSA2 | 0.60 | 0.98 |

### 3.2 Assessing Performance of PRIGSA2 on Known Protein Repeats

To show the efficacy of PRIGSA2 in accurately identifying repeat proteins, their copy number and repeat boundaries, we considered a non-redundant dataset of five subclasses of Kajava’s class III and IV repeats (listed in table 2), classified into 13 repeat families in UniProt. The non-redundant dataset of 13 families and the predicted repeat regions are given in http://bioinf.iiit.ac.in/PRIGSA2/S2_UniProt_NonRedundant.xlsx. The analysis is carried out at three levels: 1) **overall protein**: to show the ability of KPRF and *de novo* modules of the algorithm to identify known repeat proteins, 2) **copy number**: to compare the prediction overlap with UniProt annotation at the repeat copy level, and 3) **residue coverage**: to compare the prediction overlap at the amino acid level. The performance at the copy number and residue level is evaluated by computing sensitivity (TP/(TP+FN)) and precision (TP/(TP+FP)) of the proposed approach with UniProt annotation and the results are summarized in table 2. It may be noted that the performance of PRIGSA2 at the protein level is quite good for all repeat families, ∼87% for both class III and IV structural repeats. The precision values at copy number and residue coverage levels are ≥ 0.83 for 12 families (except PbH1), clearly indicating the ability of the algorithm in distinguishing between repeat and non-repeat regions in a protein. For PbH1 repeat proteins, we observed that PRIGSA2 identified larger repeat regions with more repeat copies compared to UniProt annotation. The RMSD of structural alignment of these extra copies with known PbH1 copies is <1Å indicating the accuracy of our predictions. Although the changes incorporated in PRIGSA2 algorithm lead to significant improvement in the sensitivity at both copy number and residue coverage levels compared to its previous version (see (Chakrabarty and Parekh 2014c)), we found detecting all the repeat units in HEAT protein family as HEAT challenging. We observed many instances of proteins reported as ARM in UniProt that are predicted as HEAT by PRIGSA2 and vice-versa. Because of shared ancestry, the two repeat families ARM and HEAT exhibit high similarity at both the sequence and structure level (Andrade *et al*. 2001; Gul *et al*. 2017), which is also reflected in the similarity in their *A_levc_* profiles.

**Table 2.**
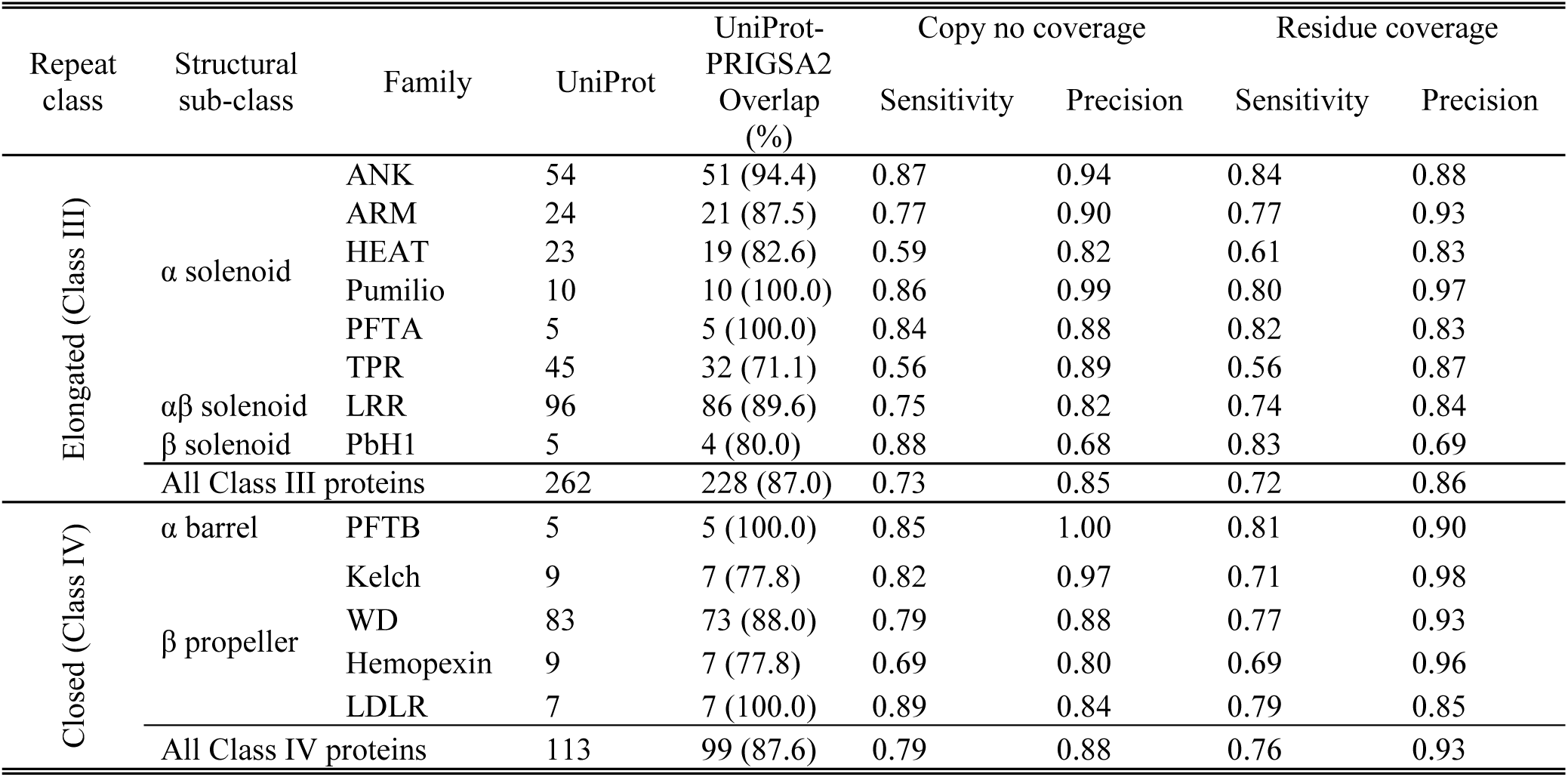
Comparison of PRIGSA2 with UniProt annotations for 13 well characterized repeat families. A non-redundant set of proteins for the 13 families is obtained by considering the best resolution structure for each UniProt entry belonging to these families. The performance at the copy number level is computed by comparing the repeat copies reported in the UniProt database with the PRIGSA2 predicted copies. The performance at the residue level shows the ability of the method to correctly identify residues belonging to repeat region.

It may be noted that the performance of PRIGSA2 on Elongated type repeat (class III) is comparable to that on Closed type repeat (class IV) indicating the ability of the program in detecting both classes of repeats equally well. However, there is still scope of improving the sensitivity of the algorithm, which is currently largely affected by secondary structure assignment programs and large insertions/deletions that affect the periodicity of peaks in the *A_levc_* profile.

### 3.3 Analysis of PRIGSA2 on PDB

On executing PRIGSA2 algorithm on all the 4,49,867 protein chains reported in the PDB (as on November 3, 2017), 11,891 repeats were predicted in 10,730 protein chains. The PRIGSA2 results on the predicted repeat proteins are given in http://bioinf.iiit.ac.in/PRIGSA2/S3_PDB_repeats.xlsx. It was observed that the membership coverage improved significantly for 12 out of 13 known repeat families (except Hemopexin) in comparison to UniProt annotation. The total coverage of these 13 families has increased by more than 3-fold from 2,500 to 8,483 PDB chains.

The *de novo* module identified 3,408 novel repeats, majority of which are uncharacterized repeat proteins. The analysis of these repeats would facilitate exploration of novel structural repeats in proteins, not yet reported in UniProt database. The predictions by *de novo* module also include members of known repeat families missed by the KPRF module or members of underrepresented repeat families in the UniProt database (< 5 proteins).

### 3.4 Analysis of de novo repeats

Below we discuss three representative examples of *de novo* predicted repeats. The first example considered is that of ‘Outer surface protein A’ 2FKJ (A) from *Borrelia burgdorferi* which forms a single layer β sheet comprising 10 copies of a structural motif (two anti-parallel β strands) of length ∼22 amino acid residues from 105 to 353 as shown in figure 7 (a). This repeat is not annotated in UniProt database (table 3), but has been classified as non-solenoid elongated repeat type (Class III.5: anti-parallel β layer/β hairpins) in Kajava’s structural classification of repeat proteins (Kajava 2012). It is reported in RepeatsDB database (Paladin *et al*. 2017) with 13 copies of ∼22 residues from 28 to 319, and is also reported as repeat protein in the literature (Makabe *et al*. 2006). The prediction is validated by performing structural alignment of all the predicted copies using Cealign module of Pymol as shown in figure 7 (d) (Shindyalov and Bourne 1998). The average RMSD of pairwise alignment between all the 10 copies predicted by PRIGSA2 is 1.4Å, indicating high confidence of prediction. Of these, 9 copies overlap with intermediate repeat copies reported in RepeatsDB. The first three N terminal copies and the last C terminal copy reported in the RepeatsDB (shown in grey color in figure 7 (a)) form a twisted curved structure different from the linear plane formed by the intermediate copies. Due to the difference in the interaction pattern in these repeat units, these units are not identified as repeat motifs by PRIGSA2 algorithm. The other example of *de novo* repeat considered is the left-handed beta helical solenoid protein acetyltransferase, 4EA9 (A) from *Caulobacter vibrioides*, shown in figure 7 (b). It contains 6 copies of tandemly repeated motif of length ∼18 residues. It is not reported in either UniProt or RepeatsDB databases, but is reported as a Hexapeptide (PF00132) repeat in literature (Thoden *et al*. 2012) and Pfam database (Finn *et al*. 2016). Three consecutive copies of Hexapeptide units, each comprising one strand and turn of equal length form an equilateral triangle structural motif arranged in a solenoid fold. The average RMSD of all-against-all pairwise structural alignment of the predicted copies is 1Å (figure 7 (e)), confirming the presence of repeat at the structure level. The third example considered is the trefoil fold repeat shown in figure 7 (c), formed by four repeat copies in Human ‘Fibroblast growth factor 1’ protein, 1HKN (chain E). It is neither reported as repeat in literature, nor in any sequence or structural repeat databases. Each repeat unit comprises two anti-parallel helices forming an overall closed structure representing the class IV repeat in Kajava’s classification. The average RMSD of all-against-all structural alignment of repeat copies in this case is 1.6Å (figure 7 (f)), confirming the prediction. These results indicate the ability of the *de novo* module of PRIGSA2 in detecting novel repeat types. Based on the length and secondary structure architecture, similar *de novo* repeat proteins can be grouped which may lead to the discovery of previously uncharacterized structural repeat families.

**Figure 7.**
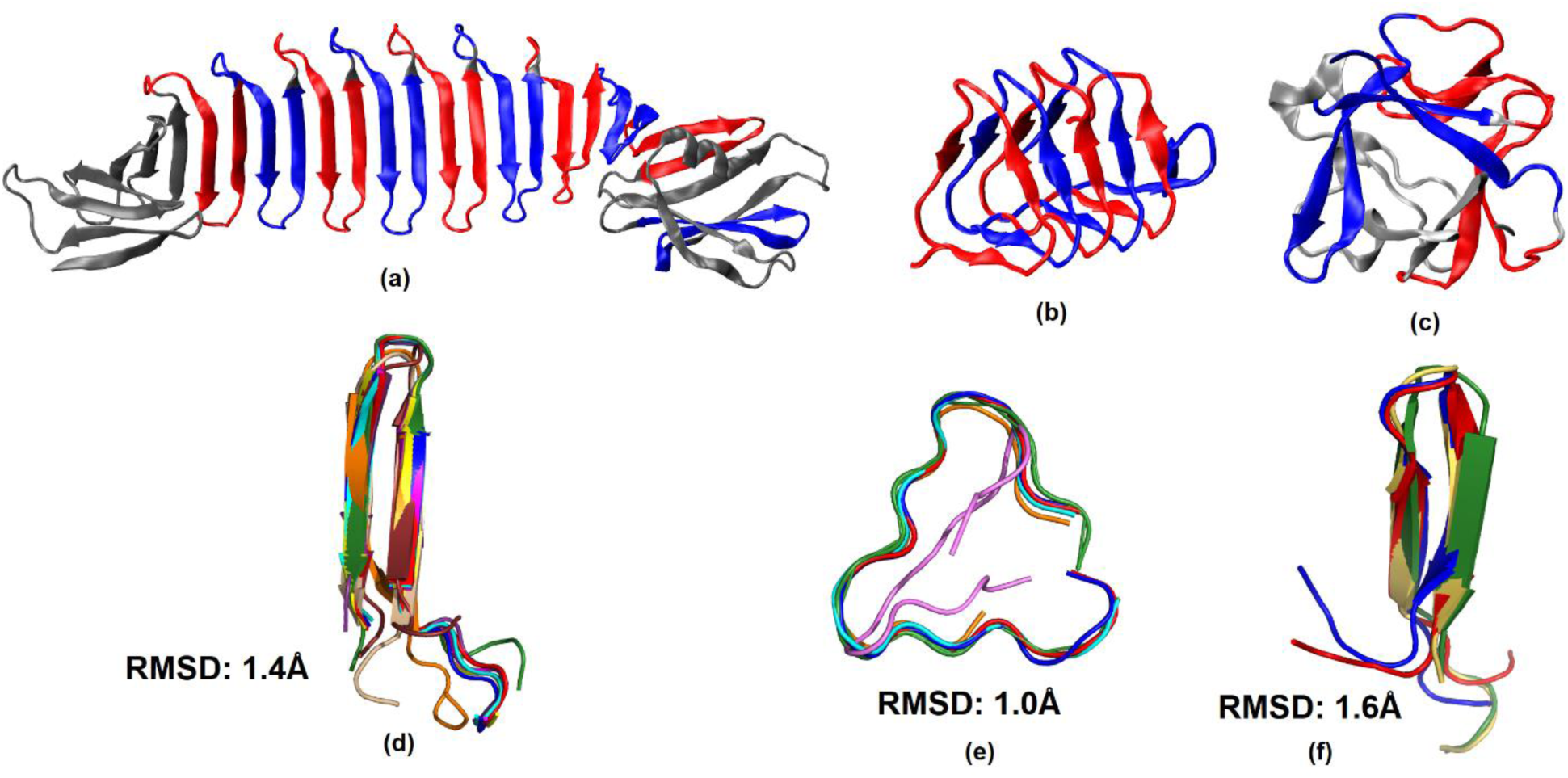
Representative examples of some repeat proteins identified by the *de novo* module of PRIGSA2 – (a) Single layer β in 2FKJ (chain A), (b) Left-handed β Solenoid in 4EA9 (chain A) and, (c) β Trefoil in 1HKN (chain E). Alternate repeat units are colored red and blue. (d), (e) and (f) show the structural alignment of all the repeat copies in (a), (b) and (c) respectively, along with the average RMSD of all pairwise alignments.

**Table 3.** Representative examples of repeat proteins predicted by *de novo* module in PRIGSA2 compared with UniProt annotation.

| PDB id (chain) | UniProt | UniProt repeat annotation | PRIGSA2 repeat annotation |
| --- | --- | --- | --- |
| 2FKJ(A) | P0CL66 | - | 22: 105-127, 128-149, 151-172, 174-195, 197-218, 220-241, 243-264, 266-287, 288-306, 333-353 |
| 4EA9(A) | O85353 | - | 18: 107-124, 126-143, 144-161, 162-178, 180-193, 195-212 |
| 1HKN(E) | P05230 | - | 21: 5018-5038, 5040-5058, 5060-5080, 5081-5099 |
| 1OBA(A) | P15057 | Cell wall-binding: 200-219, 220-239, 241-260, 261-280, 281-300, 303-322 | 20:199-213, 215-234, 236-255, 256-275, 280-295 |
| 1G9U(A) | P17778 | LRR: 72-91, 92-113, 114-131, 132-153, 154-173, 174-195, 196-215, 216-237, 238-257, 258-279, 280-297, 298-317, 318-339, 340-357, 358-379 | 40: 98-137, 140-179, 182-221, 223-263, 269-303, 309-343, 350-380 |

Some of the *de novo* predicted repeats also include members of underrepresented families in UniProt that were not considered as KPRF. For example, 1OBA is reported with 6 copies of Cell wall binding repeat type of length ∼20 residues in UniProt (table 3). Since the repeat family had less than 5 members reported in UniProt database, representative profile was not constructed in this case for the KPRF module of PRIGSA2. However, all the 5 copies of length ∼20 are identified by the *de novo* module in agreement with the annotation in UniProt as shown in table 3. Similarly, members of other underrepresented structural repeat families in UniProt such as RCC1, HAT, etc. have also been identified by the *de novo* module.

Though the overlap of PRIGSA2 with UniProt annotation is quite good for the 13 KPRFs (table 2), some members of these repeat families are missed by the KPRF module. This is either because of inaccuracy in secondary structure assignment (by STTRIDE or DSSP), poor *A_levc_* alignment, or large variations (indels) in the repeat region. For example, outer membrane protein YopM, 1G9U (chain A) is reported with 9 copies of LRR repeat in UniProt from 72 to 379 (table 3). The LRR motif comprises Strand-Helix motif, while neither DSSP nor STRIDE assignment contain any helices in the repeat region. Since the KPRF module is dependent on the conservation of consensus secondary structure architecture of a repeat family, it is not identified as LRR. Since in PRIGSA2 a structural repeat motif is defined to contain two or more secondary structural elements (any type of helix or strand), the *de novo* module predicted 7 copies of Strand-Strand repeat motif of length ∼40 residues from 98 to 380, each repeat unit covering two Strands of adjacent LRR units reported in UniProt database. Similarly, 139 other proteins that are reported as KPRFs in UniProt were detected by the *de novo* module of PRIGSA2.

### 3.5 Identifying Repeats in Multimeric Protein Complexes

In recent studies, repeat proteins have been reported that form tandem structural repeats only in *k*-meric state (Roche *et al*. 2018). The long range inter-chain interactions between the monomeric forms are responsible for the stable structure of the repeat region. To facilitate the detection of such repeats, a single network is constructed for the complete protein assembly comprising all the chains, for e.g., dimer, trimer or tetrameric protein complexes. In this case, an atom-pair contact network is constructed for the *k*-meric protein complex which is able to capture weak inter-chain interactions that are missed in the C_α_ network representation. This is illustrated for Human adenovirus C protein (PDB: 1QIU, chains: A, B, C). In figure 8 (a) is shown the overlap of *A_levc_* profiles, computed individually, for the 5 β-hairpin repeat copies in each of the three monomers using C_α_ network. It may be noted that the *A_levc_* values for the first two copies in each chain are zero, indicating that the residues in these copies have no interactions with other residues in the structure. Also, the intermediate copies do not exhibit a well-conserved pattern to be recognized by automated methods. In figure 8 (b), the overlap of *A_levc_* profiles is shown for C_α_ network constructed for the trimeric complex (i.e. a single network is constructed by considering interactions within and between the three monomers). Although the contribution of inter-chain interactions is captured to some extent, the *A_levc_* values for the first two copies of chain B are still zero and the profile of various copies is not well conserved. Figure 8 (c) depicts the overlap of *A_levc_* profiles for all the 15 copies of the trimer for atom-pair contact network representation of the trimeric complex. A good overlap in the *A_levc_* profile indicates that the interactions within and between the monomers are well captured by this network representation. Similarly, β hairpin repeats have been identified in trimeric complexes of 3S6X, 3S6Y, 3S6Z, 5N8D and 3WPA.

**Figure 8.**
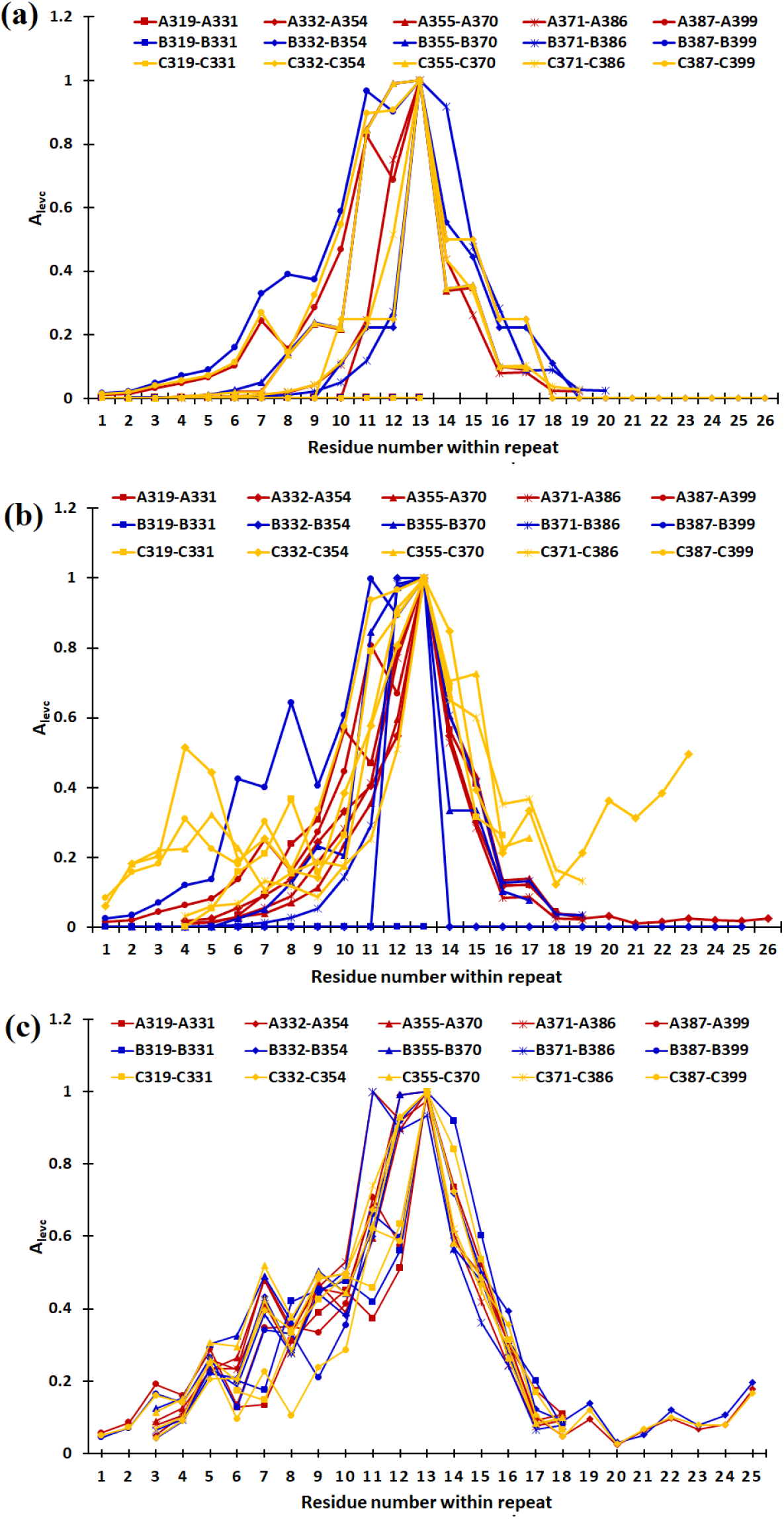
Overlap of *A_levc_* profiles of the 15 repeat copies in 1QIU chains A, B and C. (a) C_α_ network constructed for each chain separately, (b) C_α_ network constructed for the trimeric protein complex, and (c) atom-pair contact network constructed for the trimeric protein complex.

Thus, PRIGSA2 uses computationally efficient C_α_ network for the detection of repeats that form super-secondary structure fold at individual chain level, while for the detection of repeats in multimeric complexes, a more fine-grained network representation, atom pair contact network, is considered.

## 4 Conclusion

Various methods have been proposed for the detection of repeats in proteins. However, it still remains a challenging problem in protein structure analysis due to low sequence similarity between repeating units and presence of insertions and deletions within and between the repeating units. In this work we have shown that analysis of protein contact networks provide a simple and elegant approach for repeat detection at the structure level by capturing inter- and intra-repeat unit interactions in monomers as well as multimeric protein complexes. Though the performance of PRIGSA2 is comparable to other state-of-the-art algorithms, its major limitations are the dependence on correct secondary structure assignments and availability of structural data. It is observed that small secondary structure elements that are generally missed by secondary structure assignment programs, affect the prediction accuracy of PRIGSA2. The *de novo* module handles such situations by correctly predicting the repeat region, but misses out on accurate copy number detection as adjacent copies are merged (under the assumption of at least two secondary structure elements are required to form a structural motif). Though the algorithm is sensitive enough to be able to distinguish between similar repeat types, *viz.*, ARM-HEAT, ANK-TPR, Kelch-WD, etc., we believe the efficacy of the algorithm can be further improved by incorporating other structural and sequence features. On executing PRIGSA2 on the complete PDB, a large number of repeat proteins are identified by the *de novo* module. A systematic analysis of these repeat proteins can help in the identification and classification of novel structural repeats in proteins.

## Acknowledgment

BC acknowledges the support of Department of Biotechnology, Government of India for DBT BINC PhD fellowship.

## Notes

http://bioinf.iiit.ac.in/PRIGSA2/

## References

Abraham A-L, Rocha EPC, and Pothier J 2008 Swelfe: a detector of internal repeats in sequences and structures. Bioinformatics 24 1536–7

Andrade MA, Perez-Iratxeta C, and Ponting CP 2001 Protein repeats: structures, functions, and evolution. J. Struct. Biol. 134 117–31

Berman HM, Westbrook J, Feng Z, Gilliland G, Bhat TN, Weissig H, Shindyalov IN, and Bourne PE 2000 The Protein Data Bank. Nucleic Acids Res. 28 235–42

Biegert A and Söding J 2008 De novo identification of highly diverged protein repeats by probabilistic consistency. Bioinformatics 24 807–14

Chakrabarty B and Parekh N 2014a Graph Centrality Analysis of Structural Ankyrin Repeats. IJCISIM 6 305–14

Chakrabarty B and Parekh N 2014b Identifying tandem Ankyrin repeats in protein structures. BMC Bioinformatics 15 6599

Chakrabarty B and Parekh N 2014c PRIGSA: protein repeat identification by graph spectral analysis. J Bioinform Comput Biol 12 1442009

Chakrabarty B and Parekh N 2016 NAPS: Network Analysis of Protein Structures. Nucleic Acids Nucleic Acids Res. 44 W375–382

Cuff JA and Barton GJ 1999 Evaluation and improvement of multiple sequence methods for protein secondary structure prediction. Proteins 34 508–19

Do Viet P, Roche DB, and Kajava AV 2015 TAPO: A combined method for the identification of tandem repeats in protein structures. FEBS Lett. 589 2611–9

Enright AJ, Van Dongen S, and Ouzounis CA 2002 An efficient algorithm for large-scale detection of protein families. Nucleic Acids Res. 30 1575–84

Finn RD et al. 2016 The Pfam protein families database: towards a more sustainable future. Nucleic Acids Res. 44 D279–285

Frishman D and Argos P 1995 Knowledge-based protein secondary structure assignment. Proteins 23 566–79

Groves MR and Barford D 1999 Topological characteristics of helical repeat proteins. Curr. Opin. Struct. Biol. 9 383–9

Gruber M, Söding J, and Lupas AN 2005 REPPER-repeats and their periodicities in fibrous proteins. Nucleic Acids Res. 33 W239–243

Gul IS, Hulpiau P, Saeys Y, and van Roy F 2017 Metazoan evolution of the armadillo repeat superfamily. Cell. Mol. Life Sci. 74 525–41

Heger A and Holm L 2000 Rapid automatic detection and alignment of repeats in protein sequences. Proteins 41 224–37

Hirsh L, Paladin L, Piovesan D, and Tosatto SCE 2018 RepeatsDB-lite: a web server for unit annotation of tandem repeat proteins. Nucleic Acids Res. 46 W402–7

Hirsh L, Piovesan D, Paladin L, and Tosatto SCE 2016 Identification of repetitive units in protein structures with ReUPred. Amino Acids 48 1391–400

Hrabe T and Godzik A 2014 ConSole: using modularity of contact maps to locate solenoid domains in protein structures. BMC Bioinformatics 15 119

Hrabe T, Jaroszewski L, and Godzik A 2016 Revealing aperiodic aspects of solenoid proteins from sequence information. Bioinformatics 32 2776–82

Jorda J and Kajava AV 2009 T-REKS: identification of Tandem REpeats in sequences with a K-meanS based algorithm. Bioinformatics 25 2632–8

Kabsch W and Sander C 1983 Dictionary of protein secondary structure: pattern recognition of hydrogen-bonded and geometrical features. Biopolymers 22 2577–637

Kajava AV 2001 Review: Proteins with Repeated Sequence—Structural Prediction and Modeling. Journal of Structural Biology 134 132–44

Kajava AV 2012 Tandem repeats in proteins: from sequence to structure. J. Struct. Biol. 179 279–88

Main ERG, Jackson SE, and Regan L 2003 The folding and design of repeat proteins: reaching a consensus. Curr. Opin. Struct. Biol. 13 482–9

Makabe K, McElheny D, Tereshko V, Hilyard A, Gawlak G, Yan S, Koide A, and Koide S 2006 Atomic structures of peptide self-assembly mimics. Proc. Natl. Acad. Sci. U.S.A. 103 17753–8

Marsella L, Sirocco F, Trovato A, Seno F, and Tosatto SCE 2009 REPETITA: detection and discrimination of the periodicity of protein solenoid repeats by discrete Fourier transform. Bioinformatics 25 i289–295

Murray KB, Taylor WR, and Thornton JM 2004 Toward the detection and validation of repeats in protein structure. Proteins 57 365–80

Newman AM and Cooper JB 2007 XSTREAM: a practical algorithm for identification and architecture modeling of tandem repeats in protein sequences. BMC Bioinformatics 8 382

Paladin L, Hirsh L, Piovesan D, Andrade-Navarro MA, Kajava AV, and Tosatto SCE 2017 RepeatsDB 2.0: improved annotation, classification, search and visualization of repeat protein structures. Nucleic Acids Res. 45 D308–12

Patra SM and Vishveshwara S 2000 Backbone cluster identification in proteins by a graph theoretical method. Biophys. Chem. 84 13–25

Pawson T and Nash P 2003 Assembly of cell regulatory systems through protein interaction domains. Science 300 445–52

Roche DB, Viet PD, Bakulina A, Hirsh L, Tosatto SCE, and Kajava AV 2018 Classification of β-hairpin repeat proteins. J. Struct. Biol. 201 130–8

Sabarinathan R, Basu R, and Sekar K 2010 ProSTRIP: A method to find similar structural repeats in three-dimensional protein structures. Comput Biol Chem 34 126–30

Shih ESC, Gan RR, and Hwang M-J 2006 OPAAS: a web server for optimal, permuted, and other alternative alignments of protein structures. Nucleic Acids Res. 34 W95–98

Shindyalov IN and Bourne PE 1998 Protein structure alignment by incremental combinatorial extension (CE) of the optimal path. Protein Eng. 11 739–47

Sigrist CJA, de Castro E, Cerutti L, Cuche BA, Hulo N, Bridge A, Bougueleret L, and Xenarios I 2013 New and continuing developments at PROSITE. Nucleic Acids Res. 41 D344–347

Szklarczyk R and Heringa J 2004 Tracking repeats using significance and transitivity. Bioinformatics 20 Suppl 1 i311–317

The PyMOL Molecular Graphics System (Schrödinger, LLC)

Thoden JB, Reinhardt LA, Cook PD, Menden P, Cleland WW, and Holden HM 2012 Catalytic mechanism of perosamine N-acetyltransferase revealed by high-resolution X-ray crystallographic studies and kinetic analyses. Biochemistry 51 3433–44

UniProt Consortium 2015 UniProt: a hub for protein information. Nucleic Acids Res. 43 D204–212

Walsh I, Sirocco FG, Minervini G, Di Domenico T, Ferrari C, and Tosatto SCE 2012 RAPHAEL: recognition, periodicity and insertion assignment of solenoid protein structures. Bioinformatics 28 3257–64

