## Supplementary Table 1 for "PRIGSA2: Improved version of Protein Repeat Identification by Graph Spectral Analysis"

**S1 Table: The consensus *A_levc_* profile(s) and secondary structure architecture for 13 known protein repeat families used in the detection of members of these families is shown.**

| **Family** | **Pfam** | **No of UniProt entries** | **No of PDB chains** | **Number of profiles** | **Repeat length** | **SS** | **Consensus *A_levc_* profile(s)** |
| --- | --- | --- | --- | --- | --- | --- | --- |
| ANK | PF00023 | 54 | 249 | 2 | 33 | HH | 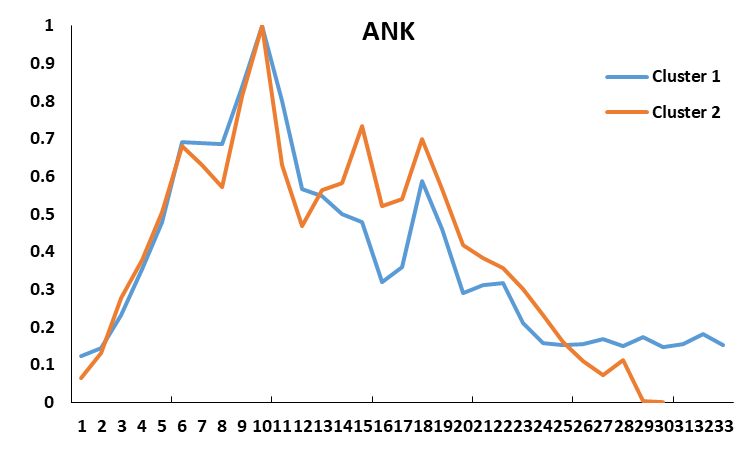 |
|  |  |  |  |  | 30 | HH |  |
| ARM | PF00514 | 24 | 230 | 2 | 42 | HHH | 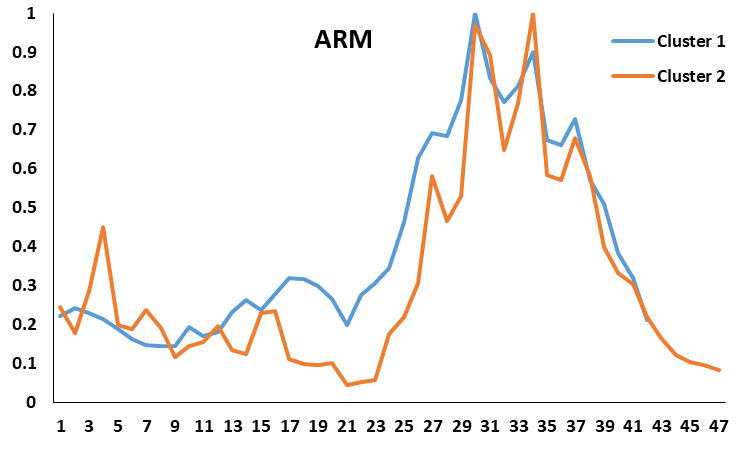 |
|  |  |  |  |  | 47 | HHH |  |
| HEAT | PF02985 | 23 | 162 | 2 | 39 | HH | 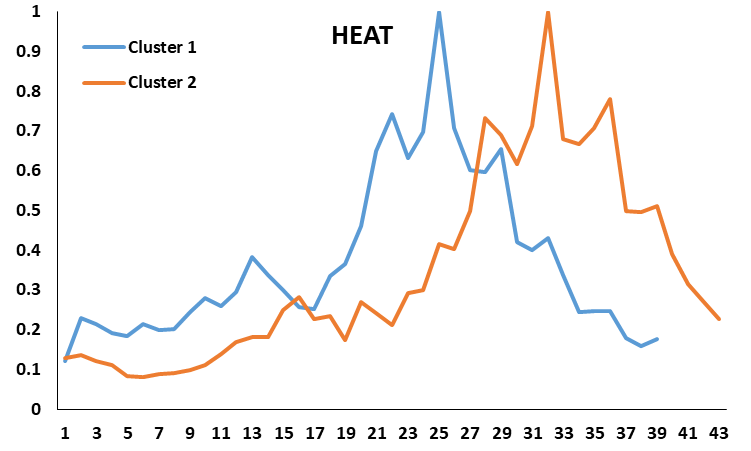 |
|  |  |  |  |  | 43 | HH |  |
| Pumilio | PF00806 | 10 | 68 | 1 | 36 | HHH | 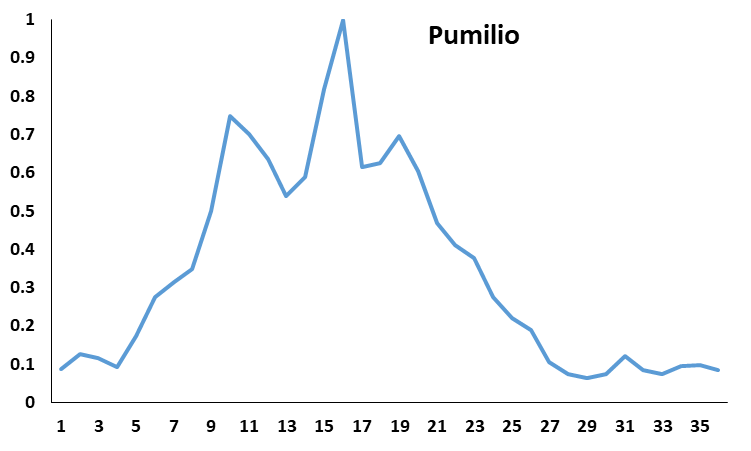 |
| PFTA | PF01239 | 5 | 144 | 1 | 35 | HH | 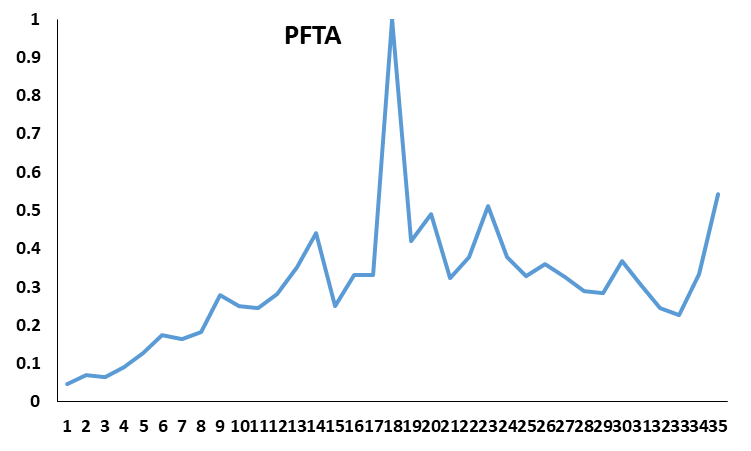 |
| PFTB | - | 5 | 79 | 2 | 42 | HH | 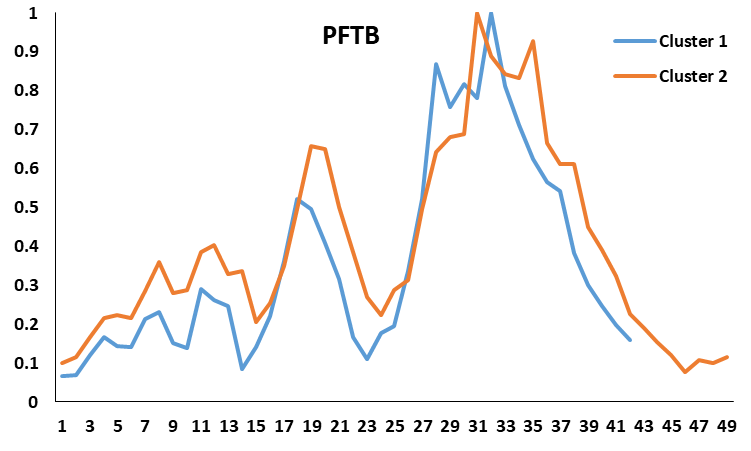 |
|  |  |  |  |  | 49 | HHH |  |
| TPR | PF00515 | 45 | 273 | 2 | 34 | HH | 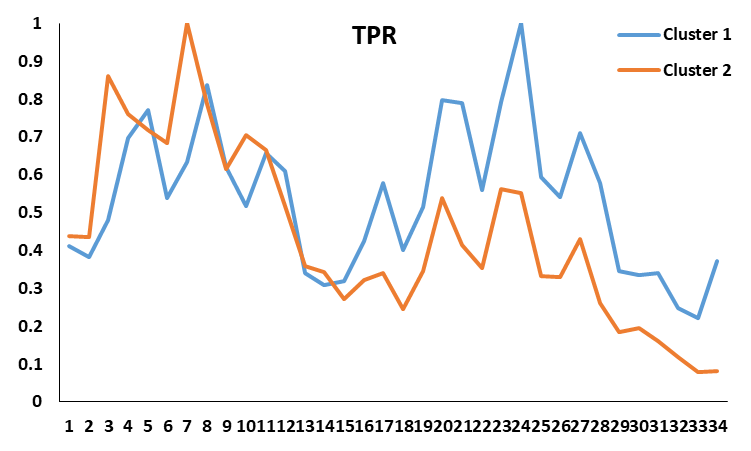 |
|  |  |  |  |  | 34 | HH |  |
| LRR | PF00560 | 96 | 525 | 2 | 22 | EH | 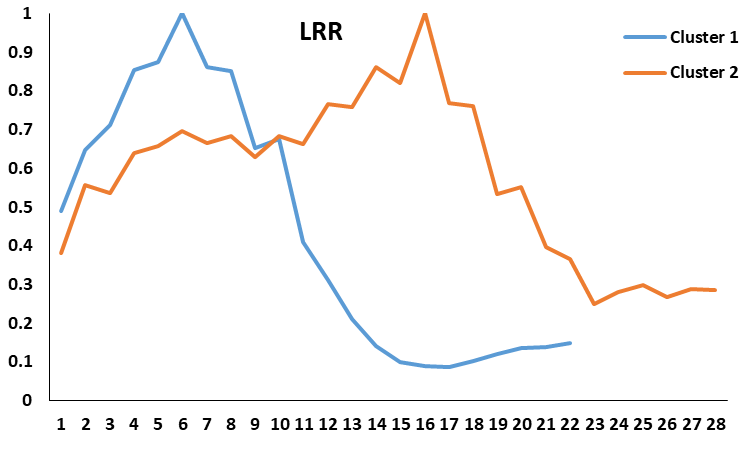 |
|  |  |  |  |  | 28 | EH |  |
| PbH1 | - | 5 | 18 | 1 | 23 | EEE | 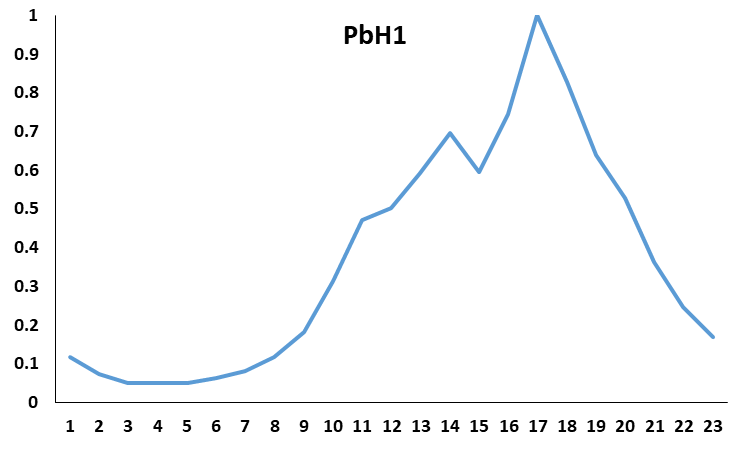 |
| Kelch | PF01344 | 11 | 67 | 1 | 47 | EEEE | 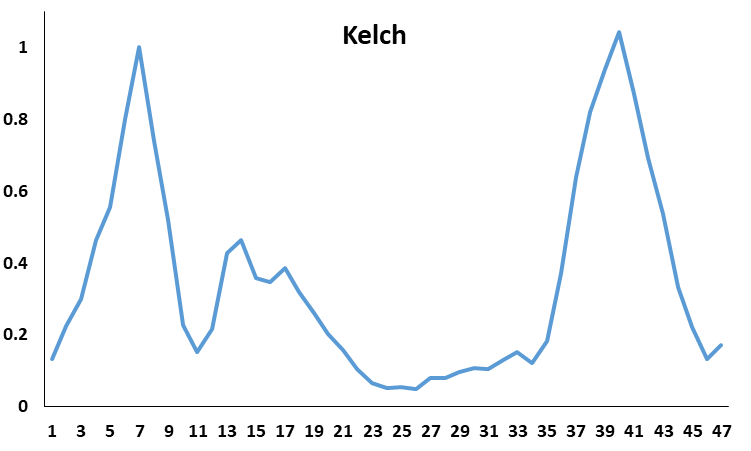 |
| WD | PF00400 | 83 | 619 | 3 | 40 | EEEE | 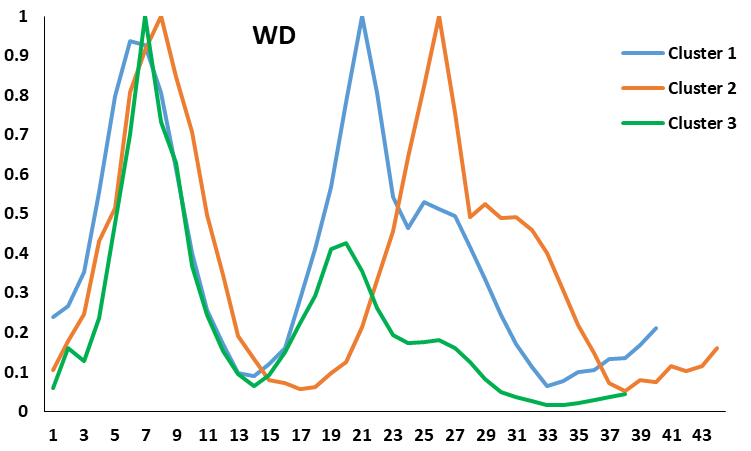 |
|  |  |  |  |  | 44 | EEEE |  |
|  |  |  |  |  | 38 | EEEE |  |
| Hemopexin | PF00045 | 9 | 35 | 2 | 49 | EEEEH | 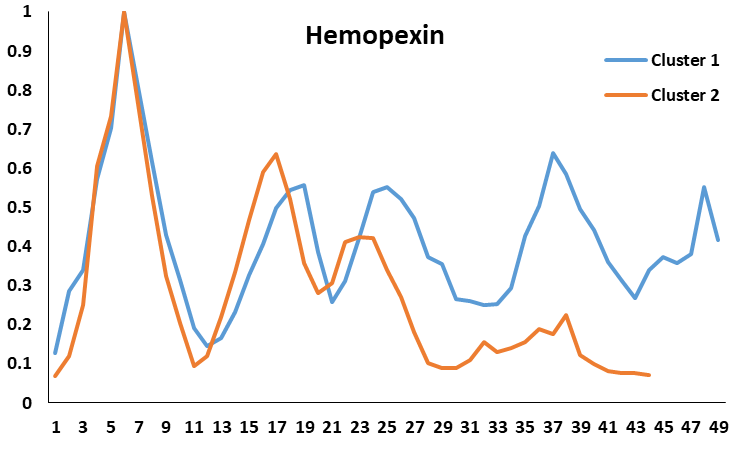 |
|  |  |  |  |  | 44 | EEEEH |  |
| LDL-receptor | PF00057 | 7 | 31 | 1 | 43 | EEEE | 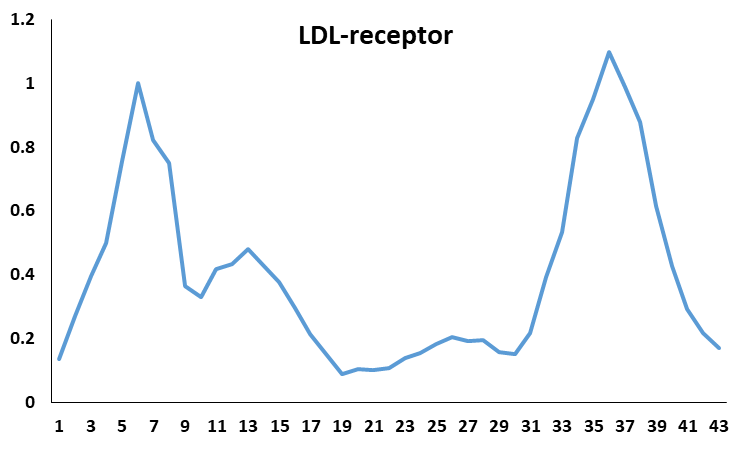 |
